# Customizable FDM-based zebrafish embryo mold for live imaging

**DOI:** 10.1101/2025.11.24.689779

**Authors:** Marcela Xiomara Rivera Pineda, Jaakko Lehtimäki, Guillaume Jacquemet

## Abstract

Accurate and reproducible orientation of zebrafish embryos is crucial for high-resolution live imaging but remains challenging with standard agarose mounting. Here, we introduce an orientation tool made using fused deposition modeling (FDM), enhanced with a thin resin coating to improve surface smoothness and performance. This design creates reproducible, embryo-shaped wells that hold larvae in a dorsoventral orientation. The molds, designed to fit standard glass-bottom imaging dishes, are inexpensive to produce on common desktop 3D printers and can be easily modified to fit different imaging setups using the open-source design files. This method enables consistent, long-term live imaging of multiple embryos and broadens the use of low-cost additive manufacturing in zebrafish imaging workflows.

## Introduction

Zebrafish (*Danio rerio*) embryos and larvae are popular vertebrate model organisms for live imaging and tracking phenomena from single-cell behavior to tissue-scale morphogenesis. This is particularly due to their optical transparency and rapid ex utero development (1). The large clutch sizes further enable simultaneous tracking and imaging of multiple age-matched embryos. The anesthetized embryos lie on their side until the inflation of the swim bladder around 120 hours post fertilization (hpf) (2). The size of the yolk further defines the position of the embryo proper, especially in younger embryos. These orientations make imaging of specific tissues, such as the brain, challenging. Therefore, immobilization techniques utilizing low-melting-point agarose or specialized molds have been developed to maintain embryos in the desired position (3–8). Due to the temperature sensitivity of embryos, agarose needs to be administered closer to the gelling point, which complicates the successful immobilization of multiple embryos in a single imaging dish for their simultaneous live imaging. Recent advances in additive manufacturing, specifically 3D printing, have enabled the creation of custom orientation tools that can produce different types of embryo-shaped wells in agarose and achieve the desired embryo orientation for imaging (4, 5, 8).

While most published 3D printing designs rely on stereolithography (SLA) for high precision (4, 5), fused deposition modeling (FDM) offers a practical, accessible alternative. FDM and SLA-based 3D printing each have their own strengths and limitations. (9) compared the two approaches in terms of performance and cost. SLA-based 3D printers can achieve very high resolution and smooth surfaces, enabling them to reproduce fine details with great accuracy. However, this comes with trade-offs, such as the need for resin handling, solvent washing, post-curing with UV light, and proper disposal of chemical waste. Conversely, FDM-based 3D printers extrude melted thermoplastic filament, such as polylactic acid (PLA). They are easier to operate and maintain, though their resolution is limited by the nozzle size and inherent material properties, resulting in less-detailed features. The two technologies also differ economically: entry-level SLA printers are generally affordable, but the ongoing costs for resin and consumables are higher. FDM printers may have a higher initial cost, but the materials are inexpensive and widely available. PLA filaments also offer a more environmentally friendly option than photopolymer resins, which produce hazardous waste. From a learning standpoint, most users find FDM easier to master, while SLA requires more attention to post-processing procedures. No single method is ideal for every situation, and both have their advantages depending on the priorities.

Here, we show that FDM is a practical, accessible method for creating molds for high-resolution live imaging of zebrafish embryos. Although it has lower print resolution and slightly less-defined cavity geometry compared to SLA, the resulting wells consistently secure larvae in a dorso-ventral position, enabling parallel, brain-focused imaging of early stages. Notably, the stability of this positioning allows immobilization with just 0.2% agarose, maintaining viability and enabling uninterrupted, high-quality time-lapse imaging for 24 hours or more, resulting in rich, cellular-level datasets over extended periods.

## Results

High-quality brain imaging of early zebrafish embryos requires positioning multiple specimens, preferably close to the coverslip surface, in a head-down orientation, which is difficult to achieve with standard agarose mounting techniques. Our first attempt at fabricating the zebrafish molds followed the approach of (10), using a 3D-printed negative to cast molds in polydimethylsiloxane (PDMS). Although these PDMS molds reproduced the embryo cavity shape more faithfully than our final fully 3D-printed versions, they presented practical challenges: the PDMS often adhered to the agarose gel, causing the gel to detach from the imaging dish. This issue led us to move toward a fully 3D-printed mold design.

Most published 3D-printed orientation molds rely on SLA printing (4, 5), but we lacked access to this technology and aimed to avoid demanding post-processing steps associated with 3D printing. Therefore, we developed an FDM-based immobilization strategy optimized for brain imaging. To design our molds, we drew inspiration from the molds proposed by (3), making some modifications to adapt them to our imaging setup. First, we introduced a circular mold format that fits standard imaging dishes with 14- and 21-mm coverslips. Both versions share the same internal slot geometry, extending fully to the glass surface. After iterative testing to improve performance, we adjusted two key slot dimensions: a longer tail and deeper slots. The tail length was increased to accommodate the size of the developmental stages we imaged, and the depth was increased to reduce surface contact between the mold and the agarose gel, helping prevent the gel from detaching from the dish. To further reduce irregularities caused by FDM printing, we applied a thin resin coating as a simple postprocessing step, which enhanced mold performance without requiring complete SLA fabrication (Fig. 1A-C). Although our molds were designed in CAD with sharp, triangular profiles, the FDM printing resolution limit of 0.4 mm nozzle size resulted in rectangular features instead (Fig. 1D). To improve handling during and after agarose gelling, we added a twopart, stamp-like design, with the upper part easily attachable or detachable when needed (Fig. 1E).

**Fig. 1.**
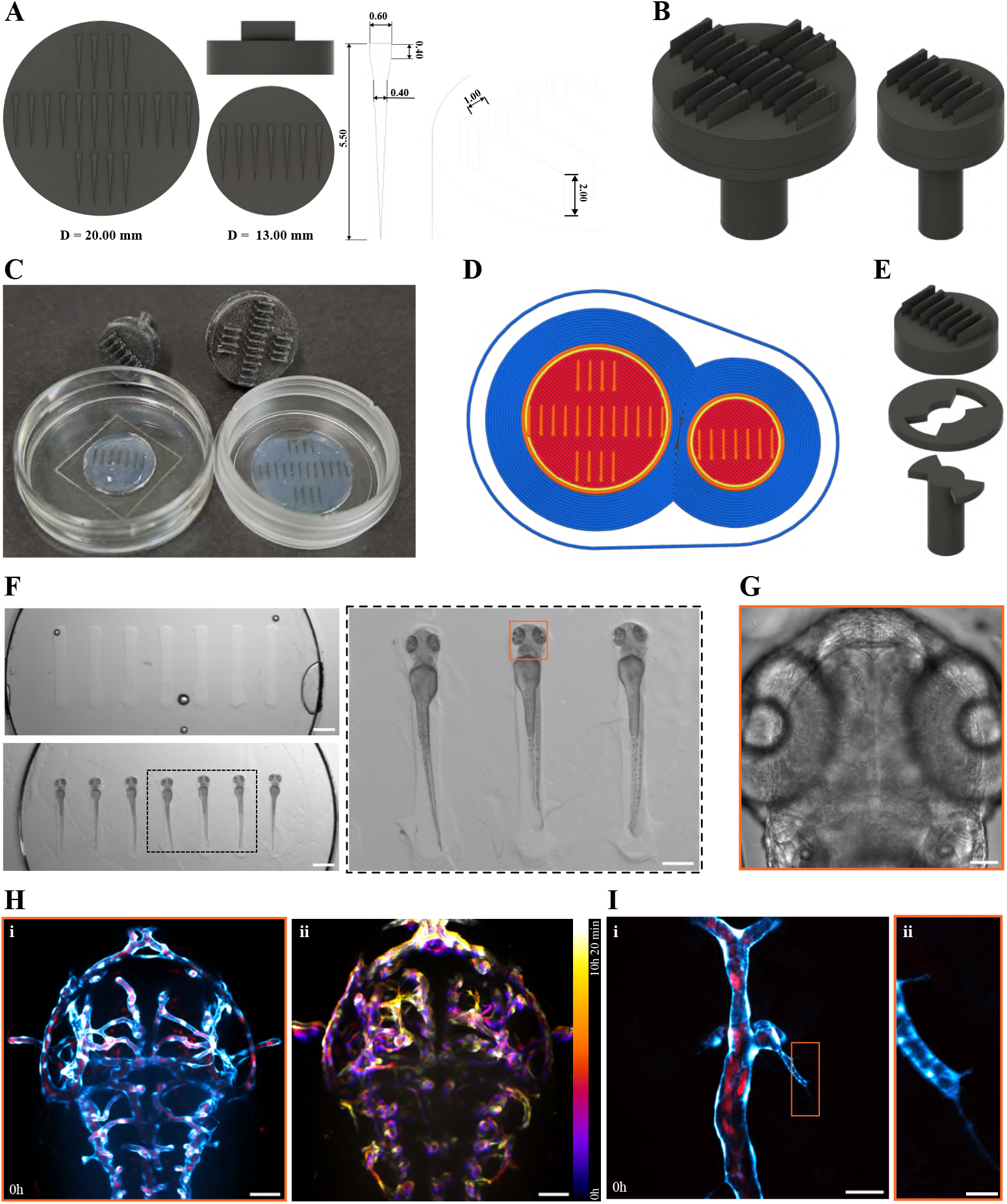
FDM-printing-based mold design improves live in vivo imaging of brain vascularization in developing zebrafish embryos. (**A**) CAD renderings of two circular mold sizes (20 mm and 13 mm) with slot dimensions in millimeters. Each slot measures 0.40 mm wide at the narrowest point, 0.60 mm at the widest, and extends 5.50 mm deep; the spacing between slots is 1.00 mm, and the slot height above the base is 2.00 mm. The smaller mold was designed for use with 14 mm coverslips, while the larger one was tested with 21 mm coverslips. The mold design was kept circular rather than rectangular to fit tightly within the well and stay level. Design files for both sizes are available for download via the associated GitHub repository. (**B**) Perspective views of the assembled molds. (**C**) Wells in 1.5% low-melting-point agarose were made using the seven- and twenty-tooth molds in 14- and 21-mm glass-bottom dishes, respectively. (**D**) PrusaSlicer g-code preview illustrating how FDM resolution (0.4 mm nozzle) converts triangular cavities into rectangular slots. (**E**) Exploded view of mold components, with a removable handle to facilitate positioning in small dishes. (**F**) Top left: image showing empty agarose wells created using the seven-teeth mold. Bottom left: image showing 56 hpf *Tg(fli1a:GFP)*;*Tg gata1a:DsRed*) embryos inserted into the wells. The black-dotted box indicates the area magnified in the next panel. Right: magnified area of mounted larvae. Live brightfield imaging of the middle larvae, marked with an orange box, is presented in panel (G). Scale bars: left panels 1 mm, right panel 500 µm. (**G**) Brightfield midplane-volume image of the 56hpf *Tg(fli1a:GFP)*;*Tg (gata1a:DsRed)* embryo from panel (F) (orange boxed region), before the start of time-lapse acquisition, shown in panels (**H-I**), orange box. Scale bar: 50 µm. (**H**) Maximum intensity projection of horizontal sections from overnight time-lapse imaging of vasculature, Tg(Fli1a:GFP, cyan), and erythrocytes, *Tg(gata1a:DsRed*, red), in the developing brain of the mounted larvae (identical to those shown in panels F-G, orange box). The first timepoint is shown in panel (i). Temporal color coding of the fli1:GFP signal over the acquisition (time-lapse, 20-minute imaging interval, 32 time points) (ii). The color bar indicates the transition from time point 1 (magenta) to 32 (bright yellow). See also **Movie S1**. Scale bars: 50 µm. (**I**) High-magnification (63x) imaging of the brain vascularization process. The maximum intensity projection of the first acquisition time point is shown in (i). The magnified area (indicated with an orange rectangle) shows filopodia-like protrusions extending from the fli1:GFP-positive (cyan) endothelial tip cell (ii). The gata1a:DsRed positive erythrocytes (red) inside a more mature blood vessel are also shown. See also **Movie S1**. Scale bars: 25 µm and 5 µm, respectively.

Using this custom orientation mold, zebrafish embryos at 50 hpf and older can be securely mounted in a dorso-ventral, head-down position, requiring only 0.2% agarose for additional immobilization (Fig. 1F-G). The mounted larvae exhibited normal development both during and after imaging when released from the agarose. The mold design prevents embryo rotation, allowing precise positioning of the heads close to the glass coverslip of the imaging dish by applying gentle downward pressure with a gel-loading pipette tip. Using *tg(fli1:GFP)* (vascular endothelial cells, cyan) and *tg(gata1a:DsRed)* (erythrocytes, red) positive embryos, this mold design enables spinning-disk confocal imaging (with 25x immersion objective) of vascular development at least up to 80 µm inside the developing brain without significant loss of signal or compromised image quality during overnight acquisition (Fig. 1H). Furthermore, as shown by the temporal projections over the acquisition period, the mold minimizes embryo movement during imaging. After an overnight session, we continued to capture detailed vascular processes from the same embryos using a high-magnification, high-NA objective and a shorter sampling interval, which allowed visualization of protrusions during vessel sprouting (Fig. 1I) with effective lateral and axial resolutions (based on a voxel size of 0.1007 x 0.1007 x 0.5 µm^3^) of approximately 0.27 and 1.0 µm, respectively. Our FDM-based mold design, enhanced with a thin resin coating and engineered for optimal brain imaging, provides a practical, reliable solution for long-term imaging of multiple zebrafish embryos from 50 hpf onward. When transferred into agarose, the cavity dimensions help position embryos in a stable dorsal-ventral, head-down orientation while requiring only a very low agarose concentration, thus preserving physiological integrity during extended liveimaging sessions. Due to their low cost, ease of fabrication, and compatibility with standard dish formats, these molds offer an accessible option for improving the consistency and quality of high-resolution zebrafish live imaging.

## Discussion

We have developed an easy and affordable method to keep zebrafish embryos properly oriented during live imaging. Using a custom FDM-printed mold to create embryo-shaped wells in agarose, this technique ensures consistent sample positioning with minimal immobilization. The mold was made from low-cost PLA filament with a resin coating and can be easily reproduced or modified; all design files and instructions are openly available through the related GitHub repository (https://github.com/CellMigrationLab/zebrafish-FDM-molds). This approach builds on previous orientation methods using 3D-printed templates (4, 5, 8) and demonstrates that despite its trade-off in precision compared to SLA printing, the FDM-printed molds remain practical and accessible tools for ensuring consistent zebrafish embryo orientation while offering a lower learning curve, reduced consumable costs, and less post-processing and handling.

In practice, the tool enables imaging the brain development of multiple larvae from a more consistent dorsal plane. Our mold design can be used from 50 hpf to 72 hpf, and likely up to 5 days post-fertilization (dpf), though not directly tested here. In brain development, these stages are commonly used to study secondary neurogenesis (11), microglial infiltration and homeostasis (12), or early neuronal activity (13). Uniform head positioning enables tissue-level features of interest, such as the midbrain vascularization process, to appear at similar spatial and axial locations across horizontal optical sections and across different imaged embryos. For this end, potential head tilting towards the side with a higher likelihood of occurring with traditional agarose mounting techniques complicates downstream visualization and analysis to separate retinal vasculature from that of the midbrain (14). The same would hold for examining other brain-resident cells, such as microglia, that inhabit both the retina and other brain regions. The two mold designs presented here can also be applied to small-scale screening experiments that commonly use 96-well plate formats with limited control over embryo orientation. Overall, and agreeing with earlier reports on standardizing embryo orientation, we find this tool improved the throughput and consistency of our imaging workflow (3–8).

The accessibility of desktop 3D printing allows most laboratories to create such tools, needing only a printer, suitable materials, and minimal post-processing. While embryos younger than 50 hpf cannot be imaged using this specific design due to the larger size of the yolk, which exceeds the diameter of the individual positions, the tooth diameter (Fig. 1A) can be easily updated in the printing setup by modifying the editable files in the related GitHub repository, enabling accommodation of younger embryos, similar to previous reports (3–5). Additionally, the tooth dimensions can be adapted for other orientations (ventral or lateral) to optimize imaging of different tissue types.

In summary, the accessibility, adaptability, and performance of the FDM-based molds offer a practical solution for achieving consistent embryo orientation and long-term live imaging, enabling standardized zebrafish mounting workflows. They also demonstrate that SLA is not the only viable option, and laboratories with existing FDM printers do not need to buy a resin printer solely for this purpose.

## Methods

### Mold design and fabrication

The zebrafish molds were modeled in the commercial computer-aided design (CAD) software Fusion 360 (Autodesk, United States). The design is a modified version of the one proposed by (3). The CAD files, in STL format, were converted to G-code using the open-source software PrusaSlicer (PrusaSlicer 2.5.0, PrusaResearch, Czech Republic). The molds were printed on a Prusa Research desktop 3D printer (Prusa i3 MK3S+, Prusa Research, Czech Republic) using polylactic acid (Prusament PLA Jet Black, Prusa Research, Czech Republic) on a textured, powder-coated print sheet (MD-28, Prusa Research, Czech Republic). All other components used the default settings. The molds were printed with the following G-code parameters: print speed of 60 mm/s, nozzle temperature of 215 °C, bed temperature of 60 °C, layer height of 0.1 mm, and infill percentage of 5 %. Each mold was designed to hold 7 and 20 embryos, respectively, from 50 hpf to 5 dpf. Printing settings are publicly available in the associated GitHub repository.

### Post-processing treatment

Printed molds were carefully sanded to remove rough edges and improve smoothness. A thin coat of two-part epoxy (Gédéo Crystal, Pébéo, 766150), mixed according to the manufacturer’s instructions, was then applied over all surfaces of the mold that contact the agarose gel. A minimum of 24 hours was allowed for the resin to cure fully.

### Zebrafish husbandry

*Tg(fli1a:GFP)* and *Tg(gata1a:DsRed)* lines were maintained and outcrossed at 28.5 °C, with embryos screened at 48 hpf (staging according to (2)) for both transgenes. Adult zebrafish were housed and used for breeding following Directive 2010/63/EU and license MMM/465/712-93 (Ministry of Forestry and Agriculture). Embryos were cultured in E3 medium supplemented with 0.2 mM 1-phenyl-2-thiourea (PTU, Fisher Scientific, 10107703) to prevent pigmentation. For embryo mounting and live imaging, E3 medium with PTU was further supplemented with 0.1 mg/ml tricaine methane sulfonate (MS-222, Sigma, E10521) to anesthetize the embryos. Live imaging was conducted on embryos between 50 and 80 hpf. Zebrafish embryos younger than 5 dpf used in experiments are not classified as protected animals under Directive 2010/63/EU.

### Agarose well preparation

High-precision 1.5 coverslips, 14 mm glass-bottom dishes (MatTek, P35G-0.170-14-C), and µ-Dish 21 mm glass-bottom dishes (Ibidi, 81158) were used. 950 µL of 1.5 % low-melting-point agarose (Sigma, A9414) in E3 medium was heated to 70 °C, then cooled to 40 °C, with 50 µL of MS-222 added and mixed thoroughly. From this mixture, 250 µL and 400 µL were added to plates with 14- and 21-mm coverslips, respectively. The molds were carefully placed on top, ensuring no air bubbles were trapped between the mold and the agarose. The agarose was allowed to solidify for 20 minutes at room temperature (RT), after which the molds were gently removed. The resulting wells (Fig. 1E) were washed with E3 medium and used immediately for embryo mounting.

### Embryo mounting

0.2% agarose was prepared by mixing 200 µl of 1 % low-melting-point agarose (at 70 °C), 750 µl of E3 medium, and 50 µl (4 mg/ml) MS-222, then incubating at 40°C. *Tg(fli1a:GFP)*;*Tg(gata1a:DsRed)* embryos at 56 hpf were anesthetized with MS-222 in E3 medium for 5 minutes. They were then added to the same E3 medium on top of the wells and gently positioned using gel loading tips (Fisher Scientific, 10411193). Excess media was aspirated, and a 0.2 % agarose solution was added dropwise to fill each well’s remaining volume. The agarose was allowed to solidify for 20 minutes at room temperature. Afterward, the plates were placed in the microscope’s imaging chamber, and 28.5 °C E3 medium with 0.1 mg/ml MS-222 and 0.2 mM PTU was gently added to cover the wells.

### Imaging

Imaging of the mounted embryos was performed using a Marianas CSU-W1 (Intelligent Imaging Innovations, 3I) spinning disk confocal equipped with a Hamamatsu sCMOS Orca Flash4.0 camera, as well as an environmental chamber and control (OkoLab), at 28.5 °C, operated via Zeiss Axio Observer 7 frame and Slidebook 6. Overnight time-lapse imaging was performed with a 25x/0.8 multi-immersion objective (LD LCI Plan-ApoC, Zeiss, 420852-9871-799) at a 20-minute interval, acquiring an 82 µm stack with a 2 µm step size. High-magnification time-lapse imaging using a 63x/1.15 water-immersion objective (LD-C-Apo, Zeiss, 421887-9970) was performed with a 2-minute interval, acquiring a 10 µm stack with a 0.5 µm step size.

### Manuscript preparation

Figures were prepared using Fiji (15) and Inkscape. GPT-5 (OpenAI) and Grammarly (Grammarly, Inc.) were employed as writing aids during manuscript preparation. The author also edited and validated all text sections. GPT-5 did not provide references. The PDF version of this manuscript was formatted using Rxiv-Maker (16).

## ABOUT THIS MANUSCRIPT

This work is licensed under CC BY 4.0.

## DATA AVAILABILITY

All design files, including technical drawings, Fusion 360 project files (.f3d), neutral CAD files (.step), and mesh files (.stl), as well as printing settings, both PrusaSlicer files (.ini and .3mf) and complete parameter configurations for reproduction in other slicers, and finalized G-code, are available in the associated GitHub repository (https://github.com/CellMigrationLab/zebrafish-FDM-molds).

## AUTHOR CONTRIBUTIONS

**Conceptualization**: J.L., M.R. **Methodology**: J.L., M.R. **Reagents**: J.L., M.R., G.J. **Formal Analysis**: J.L., M.R. **Investigation**: J.L., M.R. **Writing – Original Draft**: Everyone. **Writing – Review and Editing**: Everyone. **Visualization**: Everyone. **Supervision**: G.J. **Funding Acquisition**: J.L., G.J.

## ACKNOWLEDGEMENTS

This study was funded by the Research Council of Finland (338537 and 371287 to G.J.), the Sigrid Juselius Foundation (to G.J.), the Cancer Society of Finland (Syöpäjärjestöt; to G.J.), and the Solutions for Health strategic funding for Åbo Akademi University (to G.J.). Additionally, this research received support from theInFLAMES Flagships Programme of the Research Council of Finland (decision numbers: 337530, 337531, 357910, and 35791). This work was supported by the Research Council of Finland, FIRI 2023 grant (decision numbers: 359073, 358879) and FIRI 2024 (grant decision numbers: 367582 and 367577). Imaging was performed at the Cell Imaging and Cytometry Core, Turku Bioscience Centre, which is supported by the Finnish Advanced Microscopy Node of Euro-BioImaging Finland (Turku, Finland) and Turku Bioimaging. 3D printing was conducted at the Laboratory of Biophysics with access to facilities provided by Euro-BioImaging ERIC (Turku, Finland). The zebrafish work was conducted at the Zebrafish Core of Turku Bioscience Centre (University of Turku and Åbo Akademi University), partially supported by Biocenter Finland.

## EXTENDED AUTHOR INFORMATION

